# Domestication Compromised Microbiome-Mediated Resistance to Western Corn Rootworm in Maize

**DOI:** 10.1101/2025.10.27.684946

**Authors:** Esaú De la Vega-Camarillo, Cesar Hernández-Rodríguez, Sanjay Anthony-Babu, Julio S. Bernal

## Abstract

Western corn rootworm (WCR) (*Diabrotica virgifera virgifera*) represents a significant threat to global maize production, with annual costs exceeding $1 billion. While modern maize is highly susceptible, wild teosinte (*Zea mays* ssp. *parviglumis*) exhibits superior resistance through poorly understood mechanisms. This study investigated rhizosphere microbiome contributions to WCR resistance across the domestication gradient. We screened 23 accessions (15 wild teosinte accessions, 6 ancestral maize accessions, 2 modern maize accessions) for WCR resistance and analyzed rhizosphere microbiomes of selected resistant and susceptible accessions using Oxford Nanopore sequencing. Resistant accessions retained >80% of root structure (85.4% ± 3.2%) while supporting minimal larval survival (22.5% ± 4.8%) compared to susceptible accessions (46.7% ± 5.1% root retention, 78.6% ± 6.3% larval survival; P < 0.001). Resistant accessions recruited significantly more diverse bacterial communities under WCR pressure, with 28-31 enriched species versus 7-19 in susceptible accessions. Key enriched taxa included *Pseudomonas putida* (3.0-3.2-fold), *Stenotrophomonas maltophilia* (2.7-2.9-fold), and *Bacillus subtilis*, all possessing documented insecticidal properties. Functional analysis revealed enrichment of defense-related pathways in resistant accessions, including hydrogen cyanide production and antimicrobial compound synthesis. Wild teosinte showed the strongest responses, with significant diversity increases (P < 0.0001) and 31 enriched species under WCR herbivory. Modern maize exhibited attenuated responses regardless of resistance classification, suggesting domestication compromised plant-microbiome defensive interactions. These findings demonstrate that WCR resistance involves coordinated plant-microbiome networks and identify bacterial taxa with biocontrol potential for developing sustainable management strategies.

**Author Summary**

## INTRODUCTION

The Western Corn Rootworm (WCR) *Diabrotica virgifera virgifera* LeConte represents one of the most economically significant threats to maize (*Zea mays mays* L.) production worldwide [1,2]. This pest’s remarkable adaptability and invasive potential have earned it the designation of a “worsening pest” [3], with annual management costs and yield losses exceeding $1 billion in North America alone [2]. Traditional management strategies, primarily based on chemical pesticides and transgenic crops expressing *Bacillus thuringiensis* (Bt) proteins [4], face mounting challenges due to widespread evolution of resistance in WCR [5,6]. This concerning trend has catalyzed renewed interest in understanding and exploiting natural defense mechanisms, particularly those present in wild relatives of maize.

Teosintes (*Zea* spp. other than maize) generally, including Balsas’s teosinte (*Zea mays parviglumis* Iltis & Doebley), the wild ancestor of maize, exhibit robust constitutive and inducible defenses against herbivores, including WCR [7,8]. As such, they represent a valuable resource for understanding natural resistance mechanisms in maize. The domestication of maize from teosinte and the crop’s subsequent geographic dispersal, selection, and breeding significantly altered plant defensive traits [8,9], including changes in volatile profiles [10] and chemical defenses [7]. These evolutionary changes have potentially compromised the integrated defense mechanisms present in wild populations [8,11]. Understanding these changes and their implications for pest resistance is increasingly crucial for developing sustainable pest management strategies.

Plant resistance to herbivory extends beyond direct genetic determinants to encompass complex interactions with associated microbiota [12,13,14]. The plant microbiome, particularly in the rhizosphere, is pivotal in mediating host defense through multiple mechanisms [13,15]. These mechanisms include direct antagonism against pests, priming of plant immune responses, and modification of root chemistry [14,16,17]. Crop domestication has significantly influenced these plant-microbiome interactions [18,19], potentially affecting the recruitment and maintenance of beneficial microorganisms contributing to pest resistance.

Recent advances in understanding plant-herbivore-microbiome interactions have documented sophisticated defense mechanisms in wild plant species. For instance, certain maize varieties retain the ability to produce (E)-β-caryophyllene, a volatile compound that attracts natural enemies of root herbivores [20,21]. The rhizosphere microbiome contributes significantly to this chemical diversity [22], although modern agricultural practices may compromise these beneficial interactions [23]. Metabolomic analyses have demonstrated complex networks of defense-related compounds in both maize leaves and roots [24], with specific plant genes mediating the production of defensive volatiles [25].

The role of microorganisms in enhancing plant defense has gained increasing recognition [26], with evidence suggesting that domestication may have altered these beneficial associations [27]. However, the specific mechanisms through which the teosinte microbiome contributes to resistance against *D. virgifera* remain poorly understood. This knowledge gap represents a critical barrier to developing novel, microbiome-based strategies for sustainable pest management in maize production. Here, we investigated the microbial-mediated mechanisms contributing to Balsas teosinte’s resistance against *D. virgifera*, particularly emphasizing rhizosphere community dynamics. Through a comprehensive analysis of microbial diversity and taxonomic enrichment patterns, we identified key bacterial groups associated with enhanced plant defense, particularly among Proteobacteria. We explored their potential functional contributions through predictive metabolic analysis and correlations with plant performance under pest pressure. Our findings provide insights into the complex plant-microbe interactions potentially underlying natural pest resistance in wild teosinte and ancestral maize, contributing to the understanding of how WCR herbivory shapes rhizosphere microbial communities across the domestication gradient. Understanding these differential microbial responses between resistant and susceptible accessions may help guide future efforts to develop targeted microbial inoculants and enhance crop resilience while reducing dependence on chemical controls.

## MATERIALS AND METHODS

### Biological Materials and Growth Conditions

Balsas’s teosinte (*Zea mays* ssp. *parviglumis*) (hereafter “wild teosinte”) seeds were collected from geographic populations in Jalisco state, Mexico, following established protocols for wild maize relatives [28]. Fifteen distinct accessions were selected based on previously documented resistance characteristics [29] (S1 Table). Ancestral maize accessions included six traditional landraces representing pre-Columbian germplasm from diverse geographic regions of Mexico. Modern maize accessions consisted of two inbred lines: B73 (susceptible reference) and B73-lox9 (resistant variant with enhanced lipoxygenase activity) [30]. Before germination, all seeds were manually extracted from fruit cases using precision forceps. No surface sterilization was performed to preserve native seed-associated microbiota [31].

### Germination and Seedling Establishment

Seeds were germinated following modified protocols from Hufford et al. [32]. Briefly, seeds were placed on moistened absorbent paper towels in Petri dishes (150 × 15 mm) and maintained at 27°C for 72 hours. Individual seedlings were subsequently transferred to modified cone-tainers (4 × 25 cm, diameter × length; Stuewe and Sons, Tangent, OR, USA) following established methods for root herbivory studies [33]. Cone-tainers were modified with chiffon mesh barriers at drainage openings to prevent larval escape while maintaining soil hydraulic properties [34].

### Growth Conditions and Substrate Preparation

Plants were maintained under controlled greenhouse conditions (27 ± 1°C, 50-55% relative humidity, 16:8 light: dark photoperiod) following standardized protocols [9]. The growth substrate consisted of field soil from maize cultivation areas processed through 60-mesh sieves and homogenized with sterilized sand (1:1 v/v) as described by Robert et al. [35]. The substrate underwent three successive autoclave cycles (121°C, 103.4 kPa, 60 min) to facilitate subsequent root recovery while maintaining soil structural properties [36].

Microbiome restoration was achieved through injection with 5 mL soil tea solution (1×10^6^ CFU/mL) prepared from composite rhizosphere samples following methods adapted from Lundberg et al. [37]. Plants received measured volumes of autoclaved water at 72-hour intervals to maintain consistent soil moisture while preventing microbial contamination [38].

### Insect Preparation and Infestation Protocol Western Corn Rootworm Handling

*Diabrotica virgifera virgifera* LeConte eggs (diapause strain; Crop Characteristics, Inc., Farmington, MN, USA) were incubated following protocols described by Meihls et al. [39]. Incubation conditions were maintained at 25 ± 2°C and 80 ± 5% relative humidity for 12 ± 1 days on moistened absorbent paper. Only neonate larvae (<24 hours post-eclosion) were selected for experimental use to ensure developmental synchrony [40].

### Plant Infestation and Assessment

Following 15 days of growth, each plant was infested with 10 neonate WCR larvae using established protocols [41]. After a 10-day feeding period, plants were harvested and processed following standardized procedures for root herbivory studies [42]. Root system parameters were quantified using WinRHIZO Pro software (v.2009c, Regent Instruments Inc., Quebec, Canada), including root fresh and dry biomass, total root surface area, and volume. Aerial tissue measurements included fresh and dry biomass following standardized protocols [43].

### Larval Recovery and Processing

Larval extraction was performed using a modified Kempton recovery system as described by Kurtz et al. [44]. Recovered larvae underwent the following standardized procedures: 1. Count determination for survival assessment (larvae recovered relative to initial infestation) 2. Morphometric analysis for instar determination and instar proportion evaluation [45] 3. Fresh biomass weight measurement for larval growth assessment 4. Surface sterilization via triple rinse protocol [46] 5. Ethanol sedation and tissue homogenization for subsequent analyses.

### Selections of resistant and susceptible accessions

Accessions were classified as resistant or susceptible based on quantitative assessments of root damage, larval survival rate, and larval biomass. Classification criteria were established as follows: Root damage was assessed by measuring root area and root volume post-infestation using image analysis software. Resistant accessions were defined as those retaining >80% of their initial root area and volume, while susceptible accessions exhibited >50% reduction in root structural parameters. Larval survival rate was determined by counting the number of live larvae recovered 10 days post-infestation relative to the initial number of larvae introduced per plant. Recovery was conducted as mentioned above. Larval biomass was quantified by weighing recovered larvae individually on an analytical balance (±0.1 mg precision) after blotting dry. All measurements were conducted in triplicate for each accession, and data were subjected to analysis of variance with significance set at P < 0.05.

### Sample Collection and Processing

Rhizosphere samples were collected from two accessions (nominally resistant, susceptible) of each of the three plant groups, wild teosinte, ancestral maize, and modern maize: For teosinte these were resistant Los Naranjos, and susceptible Guachinango; for ancestral maize they were resistant Tehua and susceptible Comiteco; and for modern maize they were resistant B73-lox9 and susceptible B73. The experimental design consisted of six biological replicates per treatment, with each replicate performed three times (a total of 18 plants per variety). Plants were grown simultaneously and divided into two treatment groups: WCR-infested and non-infested controls. For the infested treatment, plants were inoculated with 10 neonate larvae per plant at day 15 after transplanting. Control plants underwent identical growing conditions but without WCR infestation. Rhizosphere samples were collected from all plants 25 days after transplanting (corresponding to 10 days post-infestation for the WCR treatment).

This temporal alignment of treatments and sampling allowed for a direct comparison of rhizosphere communities between infested and non-infested plants of the same developmental stage. Rhizosphere samples were processed within two hours of collection, with soil water content standardized to 40% WHC. When immediate processing was not feasible, samples were preserved at -80°C to maintain microbial community integrity [47].

### Microbiome Analysis

#### DNA Extraction and Sequencing

Genomic DNA extraction was performed using the ZymoBIOMICS DNA Miniprep Kit with protocol modifications specific to the sample type. For rhizosphere samples, bead beating was optimized to 6.0 m/s for 40 seconds at 4°C, followed by an additional cleanup step using 5M NaCl precipitation and double washing with Inhibitor Removal Buffer. Larval samples underwent surface sterilization through sequential washing in 1% sodium hypochlorite (1 minute), three sterile water rinses, 70% ethanol (30 seconds), and three final sterile water rinses before homogenization and DNA extraction. Final DNA elution was performed in 50 µL elution buffer with a 2-minute incubation at 70°C, followed by a second elution to maximize yield. DNA quality was assessed using NanoDrop™ spectrophotometry (acceptable A260/A280 ratios: 1.8-2.0), and quantity was determined using Qubit™ 4.0 fluorometer with dsDNA HS Assay Kit (minimum concentration requirement: 10 ng/µL) [49].

Library preparation employed the Oxford Nanopore Rapid Sequencing Kit (SQK-RAD004) with native barcoding (EXP-NBD104/114), using 400 ng of high-quality DNA per sample. Sequencing was performed on MinION R9.4.1 flow cells with a 72-hour runtime. Raw sequencing data underwent base-calling using Guppy v5.0.7 with the high-accuracy model [50], followed by adapter trimming and demultiplexing using Porechop v0.2.4. Quality control parameters included minimum read length (1000 bp), quality score (Q10), and maximum N’s percentage (1%) [51]. Initial bioinformatic processing was performed using custom Python scripts (v3.8.5) integrated with established analysis pipelines [52, 53].

### Bioinformatic and Statistical Analyses

Taxonomic and functional analyses were conducted using stringent parameters from the EZBIOME platform. Taxonomic classification employed a minimum alignment length of 1000 bp, a minimum identity threshold of 97%, and a maximum e-value of 1e-10. Alpha diversity metrics, including observed OTUs, Shannon diversity index, and Simpson’s diversity index, were calculated to capture different community richness and evenness aspects. Beta diversity was analyzed using PERMANOVA with 999 permutations and weighted UniFrac [54, 55]. Linear discriminant analysis Effect Size (LEfSe) was implemented to identify key microbial taxa distinguishing between treatment groups, with an LDA score threshold of 2.0 and a significance level of α = 0.05. This analysis focused on contrasting resistant versus susceptible varieties within each maize type and comparing responses to WCR infestation across varieties.

Functional predictions were performed using complementary approaches to ensure robust metabolic inference. FAPROTAX was employed to transform taxonomic abundance profiles into putative functional profiles based on current knowledge of microbial phenotypes [56]. Additionally, PICRUSt2 was used to predict functional gene content from marker gene data, with NSTI (Nearest Sequenced Taxon Index) scores calculated to evaluate prediction reliability [57]. The resulting functional profiles were analyzed for differential abundance using negative binomial models with FDR correction. All experimental procedures were replicated across three independent trials to ensure reproducibility and statistical robustness. Statistical significance was determined at α = 0.05, with appropriate corrections for multiple testing. Correlation analyses were performed using Spearman’s rank correlation with Benjamini-Hochberg correction [58].

## RESULTS

WCR resistance patterns in wild teosinte and ancestral and modern maize accessions revealed a complex interplay between plant resistance levels and rhizosphere microbial communities. Our comprehensive analysis showed distinct resistance profiles across modern maize, ancestral maize, and wild teosinte accessions, encompassing direct plant defenses and microbiome-mediated responses. Through integrated evaluation of WCR and plant phenotype metrics and rhizosphere microbial community analyses, we uncovered multiple defense strategies contributing to WCR resistance, particularly in wild teosinte and ancestral maize accessions.

Based on root damage assessment, resistant accessions retained >80% of their initial root area and volume (mean ± SE: 85.4% ± 3.2%), demonstrating minimal structural damage. Conversely, susceptible accessions exhibited significant reduction in root architecture, with root area and volume decreased by >50% (mean ± SE: 46.7% ± 5.1%; P < 0.001), indicating extensive larval-induced injury. Western corn rootworm larval survival differed significantly between resistance classes. Resistant accessions supported low larval survival, with mean survival rates of 22.5% ± 4.8% (range: 10-30%). In contrast, susceptible accessions maintained higher larval survival rates exceeding 70% (mean ± SE: 78.6% ± 6.3%; P < 0.001). These findings demonstrate the differential capacity of accessions to suppress larval establishment and development. Larval biomass accumulation was significantly reduced on resistant accessions, where recovered larvae averaged 1.8 ± 0.4 mg. Larvae developing on susceptible accessions achieved substantially greater biomass, averaging 7.2 ± 1.2 mg (P < 0.001), indicating enhanced nutritional quality and reduced antibiotic effects in susceptible genotypes.

We assessed WCR performance on 15 wild teosinte, 6 ancestral maize, and 2 modern maize accessions and identified groups of accessions that were most and least resistant to WCR based on evaluations of WCR and plant phenotypes (Fig. 1, 2). For example, developmental speed, while variable (Fig. 1B), was slowest in wild teosinte and ancestral maize compared to modern maize (Fig. 1A). The instar (head capsule width) distribution indicated that many WCR larvae in wild teosinte and ancestral maize remained in 1^st^-instar at the end of the experiment. At the same time, this was not evident in larvae from modern maize (Fig. 1A). Overall, most larvae recovered from wild teosinte were 1^st^-instar, followed by 2^nd^- and 3^rd^-instar; most larvae recovered from ancestral maize were 1^st^-instar and a minority were 2^nd^-instar; and most larvae recovered from modern maize were 2^nd^-instar and a minority advanced to 3^rd^-instar (Fig. 1A). This pattern suggests that domestication and modern breeding progressively reduced resistance to WCR in maize, as evidenced by the shift toward permitting development to later instars in modern maize compared to ancestral maize and wild teosinte.

**Fig. 1.**
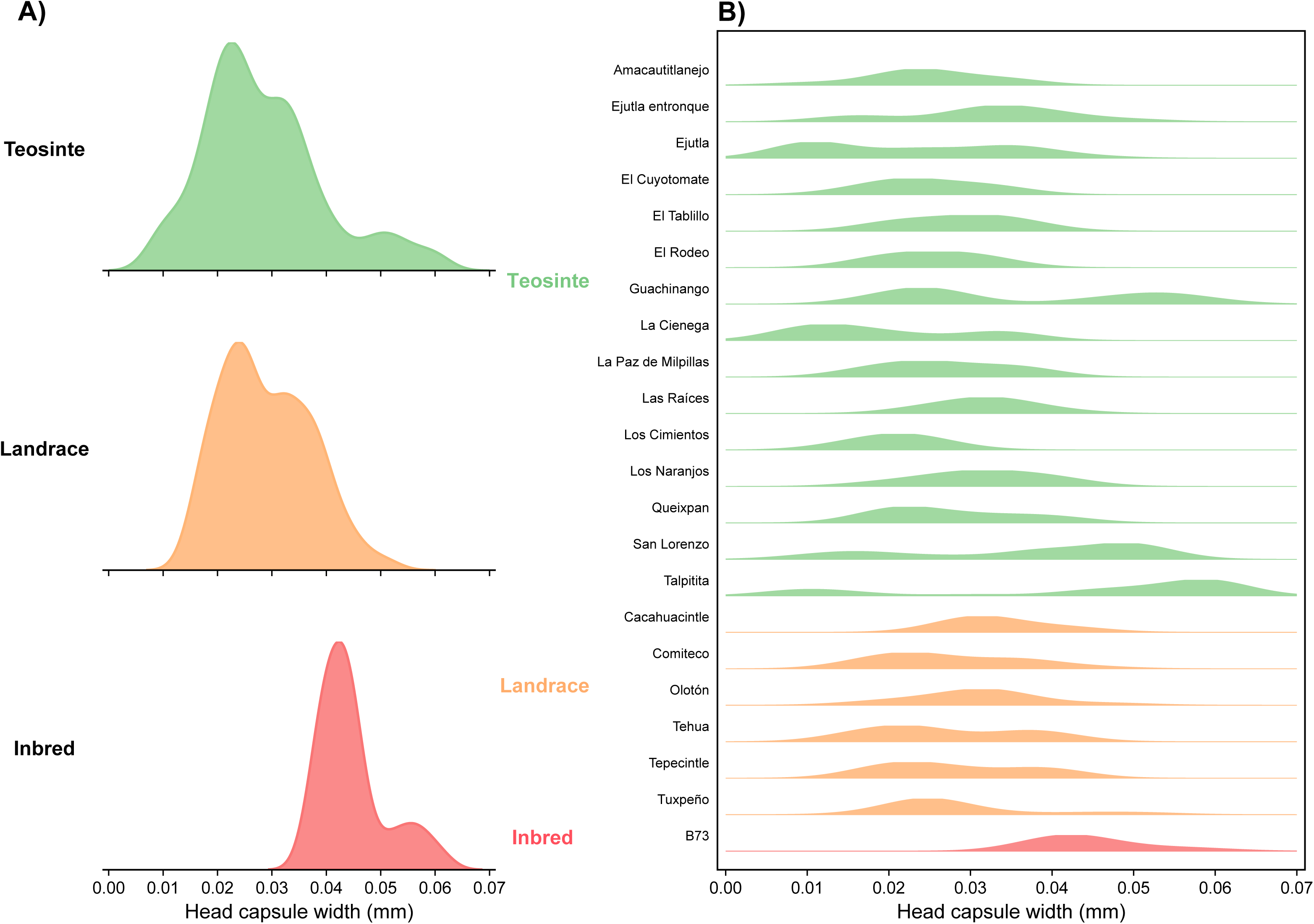
Western corn rootworm larval head capsule width distribution across plant types. Kernel density plots showing head capsule width measurements (mm) of recovered WCR larvae. (A) Average distribution across accessions within each plant type: wild teosinte (n = 15 accessions), ancestral maize (n = 6 accessions), and modern maize (n = 2 accessions). (B) Distribution including all individual accessions. Head capsule widths ≤0.06 mm correspond to 1st instar, 0.06-0.08 mm to 2nd instar, and >0.08 mm to 3rd instar larvae. Data represent measurements from larvae recovered 10 days post-infestation (n = 10 larvae per plant, 6 biological replicates per accession).

Canonical centroid analysis of three WCR performance variables (development speed, growth, survival) and two plant performance variables (belowground tolerance, aboveground tolerance) revealed resistance groups containing variable numbers of maize and teosinte accessions (Fig. 2). The first two canonical axes explained 86.15% of the total variation (PC1: 64.74%, PC2: 21.41%). The analysis separated resistant accessions (shown in green) from susceptible accessions (red), with intermediately resistant accessions (blue) occupying an intermediate position. The resistant group (green) included accessions showing strong antibiosis effects, with Los Naranjos (0.8915), Tehua (0.8461), San Lorenzo (0.8162), Queixpan Ameca (0.7722), Amacautitlanejo (0.7557), El Cuyotomate (0.7386), and Ejutla (0.7212) displaying the highest overall scores. The intermediate group (blue) included accessions La Paz de Milpillas (0.6989), La Cienega (El Fresno) (0.6822), Olotón (0.6735), El Tablillo (0.6354), and Los Cimientos (0.6143). The susceptible group (red) included accessions Ejutla entronque (0.5810), Talpitita (0.5492), El Rodeo (0.4939), Cacahuacintle (0.4850), Tepecintle (0.4355), Tuxpeño (0.4355), Las Raices (0.4290), Guachinango (0.3189), Comiteco (0.1126), and B73 (0.0218). It showed positive correlations with larval performance vectors, indicating favorable conditions for WCR development. Statistical comparisons revealed significant differentiation between the resistant and tolerant groups for biomass reduction (p=0.0335), average larvae weight (p = 0.0248), and biomass infested (p = 0.0039). Resistant vs. susceptible groups comparisons showed significant differences in biomass reduction (p = 0.0020), average larvae weight (p = 0.0177), and average survival (p = 0.0008). In contrast, intermediate vs susceptible groups differed significantly in biomass infested (p = 0.0020) and average survival (p = 0.0079). This multivariate pattern suggested the existence of multiple, distinct resistance mechanisms operating in teosinte accessions compared to maize.

**Fig. 2.**
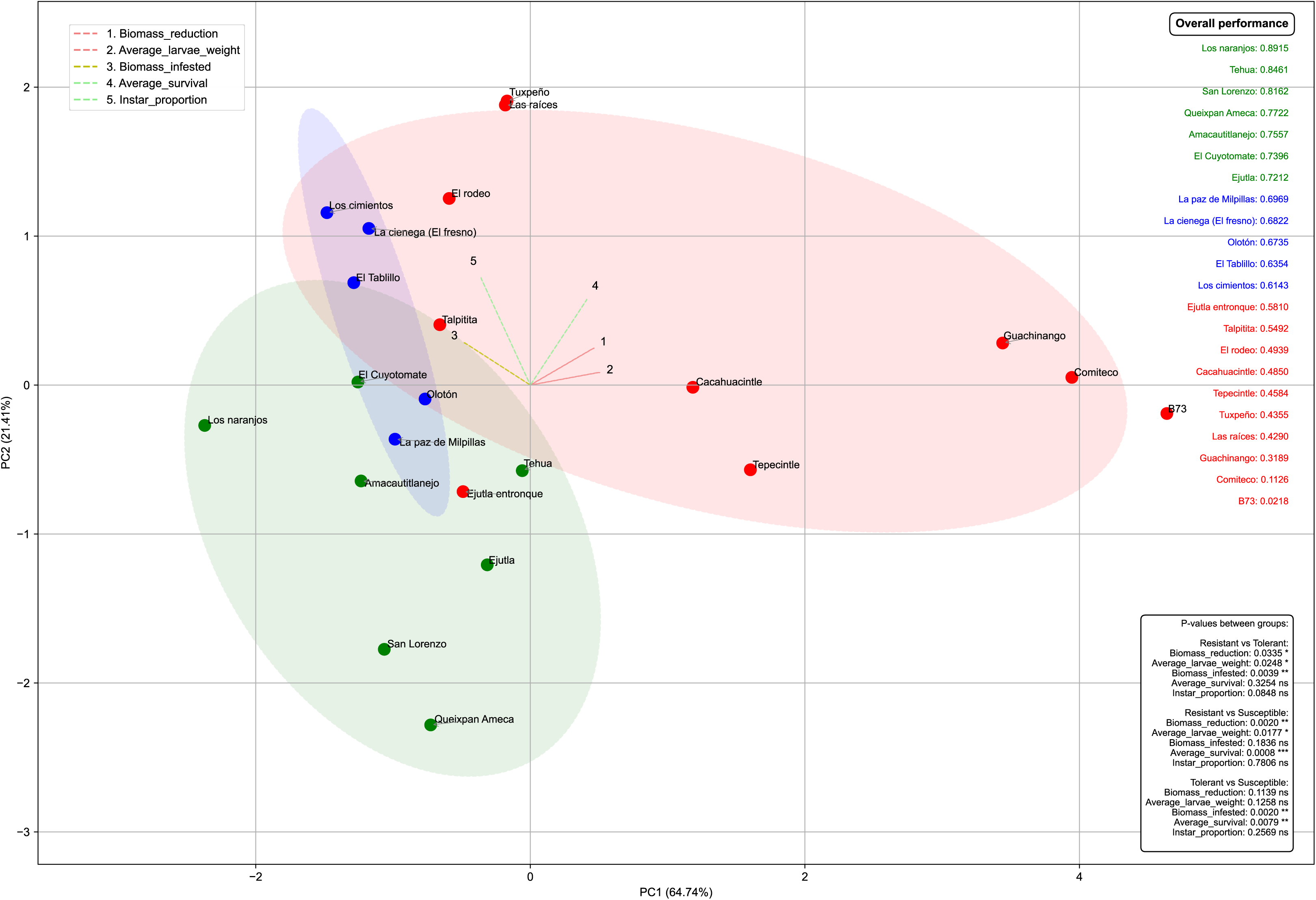
Canonical centroid analysis of WCR resistance metrics. Principal component analysis based on five performance variables: WCR development speed, larval growth, larval survival, plant belowground tolerance, and plant aboveground tolerance. PC1 explains 64.74% of variance, PC2 explains 21.41% (total: 86.15%). Resistant accessions (green circles, n = 7): Los Naranjos (0.8915), Tehua (0.8461), San Lorenzo (0.8162), Queixpan Ameca (0.7722), Amacautitlanejo (0.7557), El Cuyotomate (0.7386), Ejutla (0.7212). Intermediate accessions (blue circles, n = 5): scores 0.6143-0.6989. Susceptible accessions (red circles, n = 10): scores 0.0218-0.5810. ANOVA with Tukey’s HSD post-hoc test, significance at p < 0.05.

Alpha diversity analysis of rhizosphere microbial communities revealed distinct patterns between resistant and susceptible genotypes across maize genetic backgrounds when exposed to WCR larvae (Fig. 3). We evaluated three diversity metrics: richness of Operational Taxonomic Units (OTUs), Phylogenetic Diversity, and Shannon’s diversity index. While OTU richness and phylogenetic diversity increased with exposure to WCR in both the resistant and susceptible wild teosinte accessions, the increases in the former (p<0.0001 for both diversity metrics) were stronger than in the latter (p = 0.003, p = 0.012). Interestingly, Shannon’s diversity index increased with exposure to WCR in the resistant wild teosinte accession (p<0.0001) but not in the susceptible accession (ns, p≥0.017). In contrast to wild teosinte, the responses of the resistant and susceptible accessions of both ancestral maize and modern were consistently similar in each of the three-diversity metrics (p≤0.005); additionally, the diversity metrics did not differ significantly in the absence and presence of WCR (ns, p≤0.017) (Fig. 3). Overall, these results suggested that resistant and susceptible teosinte accessions show divergent responses to WCR herbivory, while resistant and susceptible maize accessions do not show divergent responses.

**Fig. 3.**
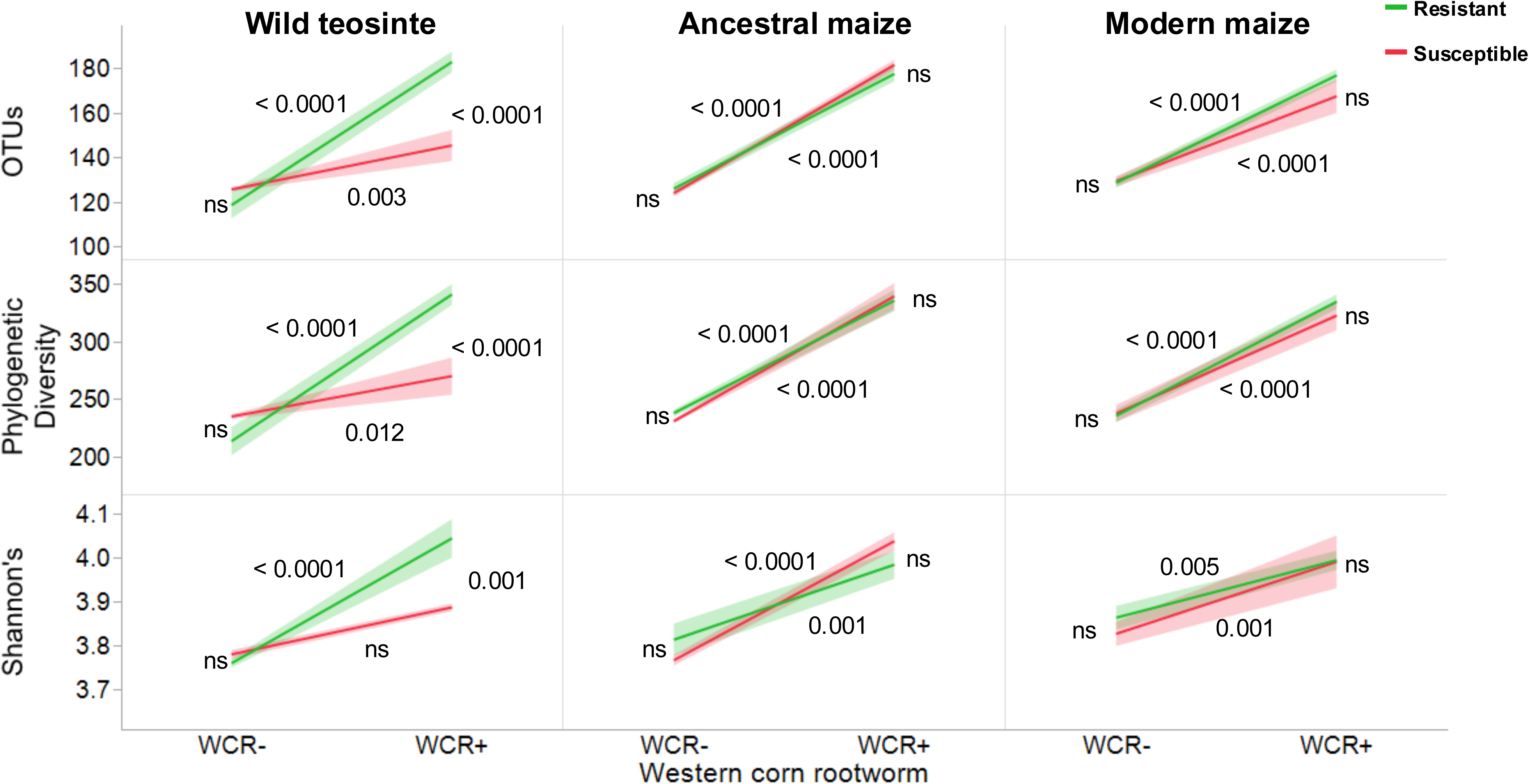
Alpha diversity metrics of rhizosphere bacterial communities. Three diversity indices measured in resistant and susceptible accessions under control (WCR-) and WCR-infested (WCR+) conditions. Top panels: OTU richness (number of operational taxonomic units). Middle panels: Phylogenetic diversity index. Bottom panels: Shannon’s diversity index. Wild teosinte resistant: OTU richness increased from ∼125 to ∼180 (p < 0.0001), phylogenetic diversity from ∼220 to ∼340 (p < 0.0001), Shannon’s from ∼3.75 to ∼4.05 (p < 0.0001). Wild teosinte susceptible: smaller increases with p = 0.003, 0.012, and ns respectively. Ancestral and modern maize showed consistent responses between resistant and susceptible accessions (all p ≤ 0.005). Data from n = 6 biological replicates per treatment. Statistical analysis: paired t-tests with Benjamini-Hochberg correction.

Multivariate analysis of rhizosphere bacterial communities revealed distinct compositional patterns between control and WCR-infested plants across the accessions. Principal Coordinate Analysis based on weighted UniFrac distances demonstrated consistent separation along PC1, which explained 84.61% of the total variation (Fig. 4A-C). In the accession × WCR (control, treated) visualization (Fig. 4A, D), all six combinations maintained their identities while showing clear separation by WCR treatment status. The resistance (resistant, susceptible) × accession visualization (Fig. 4B, E) revealed clustering driven primarily by WCR treatment and resistance categorization within WCR treatment. Finally, the WCR treatment-centered visualization (Fig. 4C, F) showed the importance of WCR herbivory in driving the plant microbiome community. Most clearly, the importance of WCR herbivory in mediating the composition of the plant microbiome in the UPGMA dendrograms (Fig. 4D-F). Altogether, these results provided robust evidence that WCR herbivory is the principal determinant of rhizosphere bacterial community structure, superseding variation associated with plant accession and WCR resistance categorization.

**Fig. 4.**
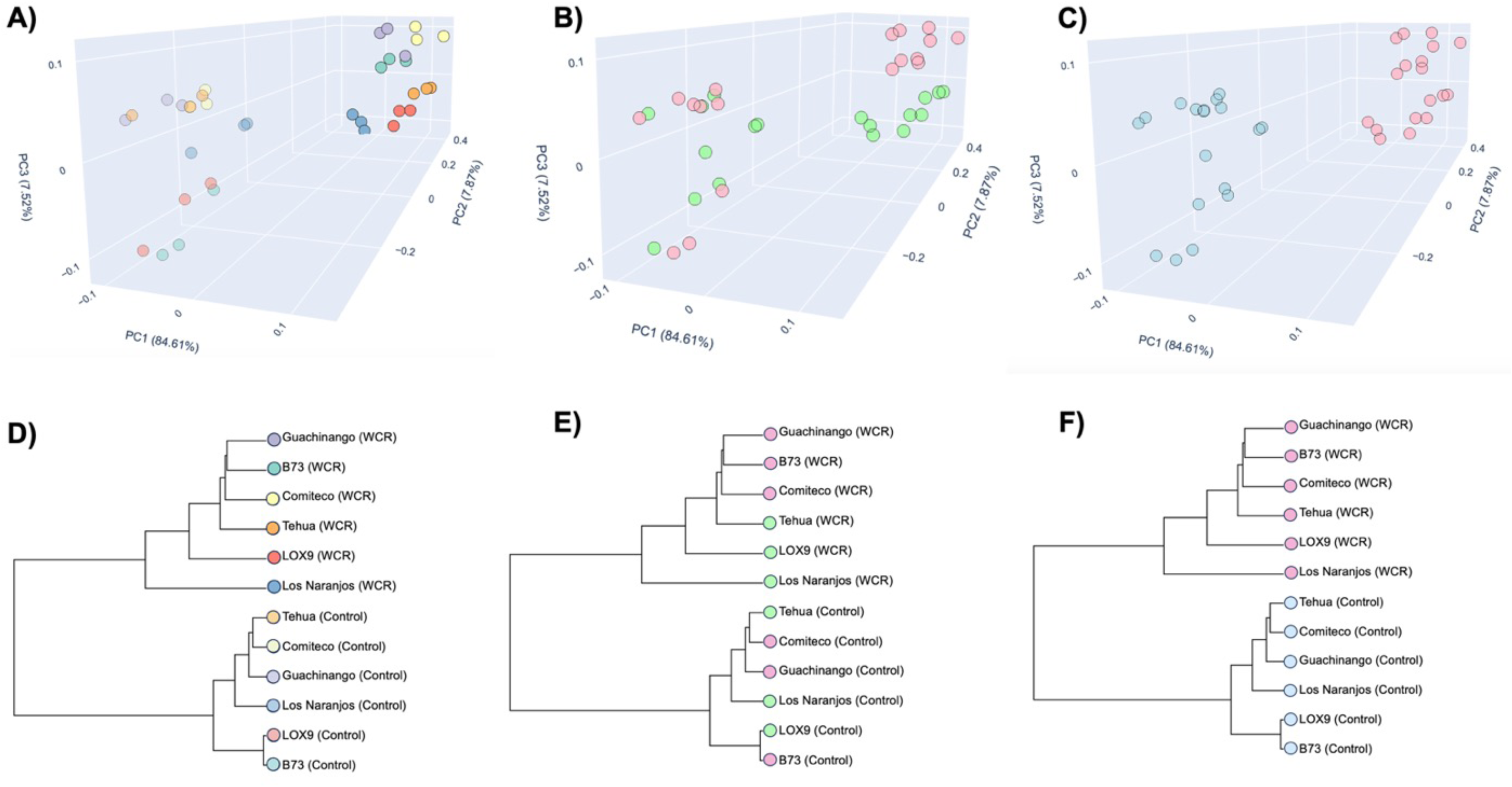
Beta diversity analysis using weighted UniFrac distances. Principal Coordinate Analysis showing rhizosphere bacterial community composition. (A) Samples colored by accession × WCR treatment combination. (B) Samples colored by resistance category × accession type. (C) Samples colored by WCR treatment status. PC1 explains 84.61% of total variation. (D-F) Corresponding UPGMA dendrograms based on weighted UniFrac distances showing hierarchical clustering patterns. Analysis based on Oxford Nanopore 16S rRNA gene sequences (minimum 1000 bp, 97% identity threshold, e-value 1e-10). PERMANOVA with 999 permutations used to test significance of group separations. n = 6 biological replicates per accession per treatment.

To examine the directional shifts in rhizosphere microbial communities following WCR infestation, we tracked community transitions across the resistant and susceptible accession groups in all three plant groups (Fig. 5). The diagram shows inconsistent directional changes in community composition (arrows connecting the control and WCR-infested states for each plant group × resistance category combination) with modern maize (both resistant and susceptible) changing in one direction (downward) and ancestral maize and wild teosinte changing in the opposite direction. Overall, these results suggested that WCR resistance involves multiple defense strategies that have been progressively eroded through domestication and modern breeding. Wild teosinte and ancestral maize accessions exhibited superior resistance through both direct antibiosis effects and differential rhizosphere microbial community responses. In contrast, modern maize showed increased susceptibility and uniform microbial responses regardless of resistance classification.

**Fig. 5.**
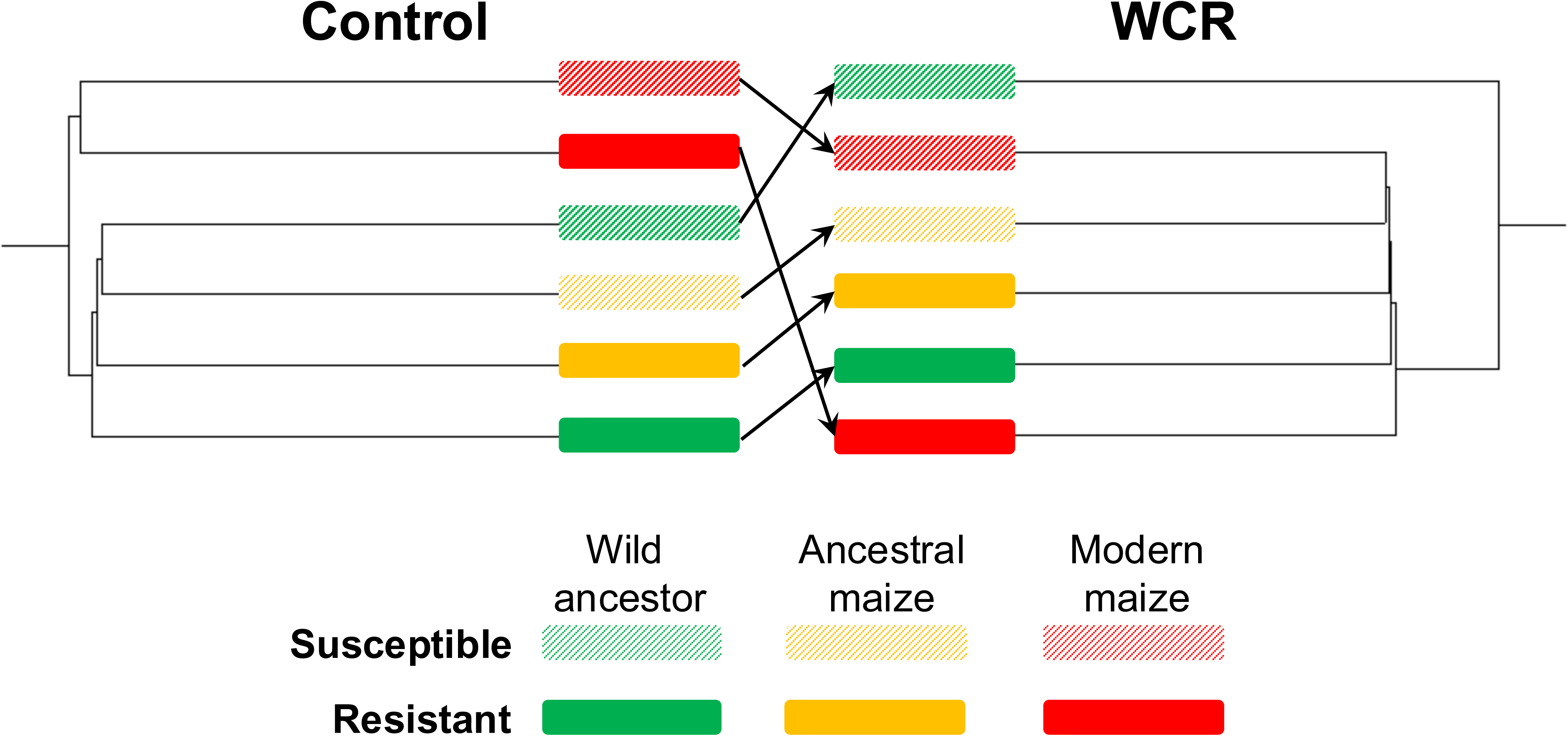
Community composition transitions under WCR herbivory. Alluvial diagram showing directional changes in rhizosphere microbial community composition from control to WCR-infested states. Each horizontal bar represents community composition for one accession-resistance combination. Arrows connect control and WCR treatments for the same accession. Wild ancestor (teosinte): resistant (green solid) and susceptible (green hatched). Ancestral maize: resistant (yellow solid) and susceptible (yellow hatched). Modern maize: resistant (red solid) and susceptible (red hatched). Community composition based on weighted UniFrac distances from bacterial 16S rRNA gene sequencing data.

A detailed examination of rhizosphere microbial communities has evidenced distinct response patterns across accessions and WCR resistance categories (Fig. 6). The resistant ancestral maize showed 46 bacterial species in the absence of WCR, of which 29 species were enriched with WCR herbivory. Notable enrichments included members of Proteobacteria (*Pantoea dispersa*: 3.2-fold, *Pseudacidovorax intermedius*: 3.0-fold), followed by *Herbaspirillum huttiense* (2.8-fold) and the *Stenotrophomonas maltophilia* group (2.7-fold). The Enterobacterales order represented 22% of the enriched species. The resistant wild teosinte exhibited 47 species in the absence of WCR, of which 31 species were enriched with WCR herbivory. Enrichment was observed in *Pseudomonas putida* (3.0-fold), *Stenotrophomonas maltophilia* (2.8-fold), and *Enterobacter tabaci* (2.5-fold). Notably, beneficial soil bacteria, including *Bacillus subtilis* and *Rhizobium* species, comprised 18% of the enriched community. The resistant modern maize harbored 49 species, of which 28 were enriched with WCR herbivory. The enriched species included *Pseudomonas putida* (3.2-fold), *Stenotrophomonas maltophilia* (2.9-fold), and multiple *Enterobacter* species (average 2.7-fold). *Herbaspirillum* and *Mitsuaria* species showed consistent enrichment patterns. Among susceptible accessions, the susceptible modern maize hosted 45 species, of which 16 were enriched with WCR herbivory; susceptible ancestral maize hosted 48 species, of which 19 were enriched, predominantly *Pantoea dispersa* (3.5-fold) and *Herbaspirillum huttiense* (3.2-fold); and susceptible wild teosinte hosted 30 species of which only seven were enriched, including *Stenotrophomonas* (2.1-fold) and *Acinetobacter lwoffii* (2.0-fold) (Fig. 6). Overall, these results suggested that resistant accessions recruit substantially more diverse bacterial communities in response to WCR herbivory compared to susceptible accessions, with wild teosinte showing the most restricted enrichment patterns.

**Fig. 6.**
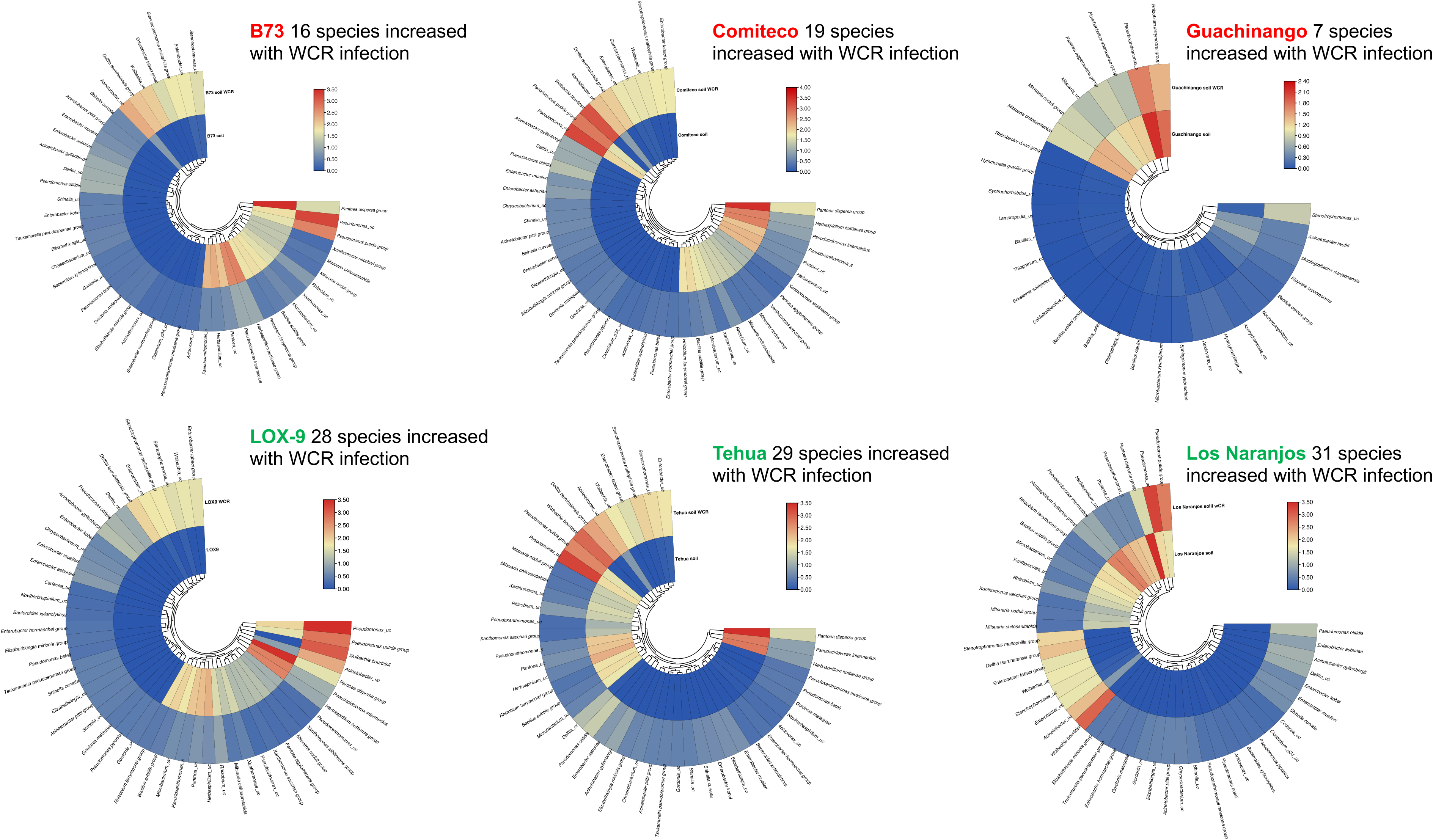
Bacterial species fold-change enrichment under WCR herbivory. Log2 fold-change values for bacterial species showing significant enrichment (≥2.0-fold increase) in WCR-treated vs. control rhizosphere samples. Resistant accessions: Los Naranjos (31 enriched species), Tehua (29 species), LOX9 (28 species). Susceptible accessions: Comiteco (19 species), B73 (16 species), Guachinango (7 species). Top enrichments include *Pseudomonas putida* group (3.0-3.2-fold), *Stenotrophomonas maltophilia* group (2.7-2.9-fold), *Pantoea dispersa* group (3.2-3.5-fold). Statistical significance determined by DESeq2 negative binomial models with FDR correction (q < 0.05). Taxonomic classification: minimum 1000 bp alignment, 97% identity threshold.

We used complementary functional prediction approaches (FAPROTAX and PICRUSt2) to identify paired comparisons between control and herbivory conditions in which putative metabolic functions were significantly enhanced or enriched under WCR herbivory across resistant and susceptible accessions. Five such comparisons were found to be significant following FDR correction (Fig.7). In resistant wild teosinte, several bacterial functions showed significant enrichment with WCR herbivory, including antibiotic production (log2FC = 3.2, -log10 p = 1.8), hydrogen cyanide production (log2FC = 2.6, -log10 p = 1.5), DAPG synthesis (log2FC = 2.8, -log10 p = 1.7), benzoxazinoid metabolism (log2FC = 2.9, -log10 p = 1.6), and chitinase production (log2FC = 2.4, -log10 p = 1.4) (Fig. 7A). This pattern suggested a coordinated microbiome response to insect herbivory in the maize wild ancestor. In wild teosinte under WCR herbivory, the resistant versus susceptible teosinte comparison revealed significant differences in microbial functional profiles (Fig. 7B). The enriched functions in resistant teosinte included quorum sensing regulation (log2FC = 3.5, -log10 p = 1.6), pyoverdine synthesis (log2FC = 2.7, -log10 p = 1.4), and protease inhibitors (log2FC = 2.5, -log10 p = 1.4). Depleted functions included lipopeptides (log2FC = -2.7, -log10 p = 1.4). This suggested distinct bacterial community functions related to defense against WCR herbivory.

**Fig. 7.**
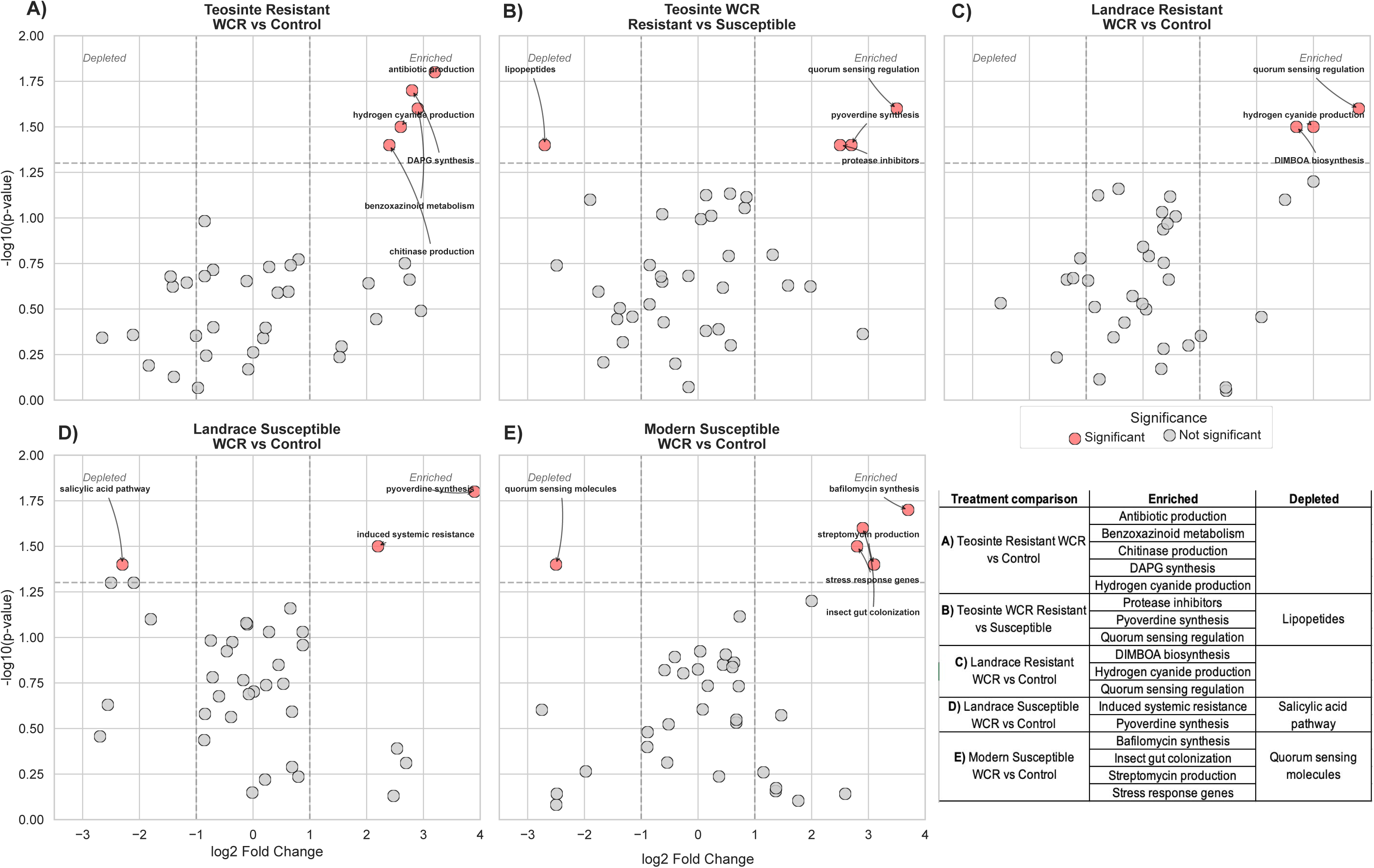
Functional pathway enrichment analysis. Volcano plots showing differential abundance of predicted microbial functions. X-axis: log2 fold-change (WCR vs. control). Y-axis: -log10 p-value. Red circles: significantly enriched functions (FDR q < 0.05, |log2FC| > 1.0). (A) Resistant wild teosinte: antibiotic production (log2FC = 3.2, -log10 p = 1.8), hydrogen cyanide production (log2FC = 2.6, -log10 p = 1.5), DAPG synthesis (log2FC = 2.8, -log10 p = 1.7). (B) Resistant vs. susceptible wild teosinte under WCR: quorum sensing regulation (log2FC = 3.5, -log10 p = 1.6). (C) Resistant ancestral maize: quorum sensing regulation (log2FC = 3.8, -log10 p = 1.6), DIMBOA biosynthesis (log2FC = 2.7, -log10 p = 1.5). (D) Susceptible ancestral maize: pyoverdine synthesis (log2FC = 3.9, -log10 p = 1.8). (E) Susceptible modern maize: bafilomycin synthesis (log2FC = 3.7, -log10 p = 1.7). (F) Summary table of enriched/depleted functions. Functional predictions: FAPROTAX and PICRUSt2 with NSTI quality scores.

Significant differences were found in comparisons between WCR-treated plants and controls in both the resistant and susceptible ancestral maize. Distinct microbiome functional changes were evident in resistant ancestral maize (Fig. 7C). The enriched functions were primarily related to antimicrobial compounds, including quorum sensing regulation (log2FC = 3.8, -log10 p = 1.6), hydrogen cyanide production (log2FC = 3.0, -log10 p = 1.5), and DIMBOA biosynthesis (log2FC = 2.7, -log10 p = 1.5). These benzoxazinoid-related pathways suggest activation of plant defensive chemistry in the resistant ancestral maize. A divergent pattern of functional enrichment was evident in the case of susceptible ancestral maize (Fig. 7D). In this case, the enriched pathways included pyoverdine synthesis (log2FC = 3.9, -log10 p = 1.8) and induced systemic resistance (log2FC = 2.2, -log10 p = 1.5), and depleted functions included the salicylic acid pathway (log2FC = -2.3, -log10 p = 1.4). This suggested potential suppression of plant defense signaling in the susceptible ancestral maize.

Bacterial functions showed significant alterations in the susceptible modern maize (Fig. 7E). The significantly enriched functions included bafilomycin synthesis (log2FC = 3.7, -log10 p = 1.7), streptomycin production (log2FC = 3.1, -log10 p = 1.6), stress response genes (log2FC = 2.8, -log10 p = 1.5), and insect gut colonization (log2FC = 2.9, -log10 p = 1.6). Significantly depleted functions included quorum-sensing molecules (log2FC = -2.5, -log10 p = 1.4). These results suggested that WCR herbivory triggers distinct metabolic responses in the rhizosphere microbiome depending on plant resistance status and genetic background. Resistant ancestral maize appeared to recruit microbiomes capable of supporting plant defensive chemistry, particularly benzoxazinoid biosynthesis and antimicrobial compound production. In contrast, susceptible ancestral maize showed evidence of compromised defense signaling, with depletion of salicylic acid pathways potentially facilitating WCR establishment.

Altogether, these patterns (Fig. 7F) collectively suggested a progressive loss of defensive microbial functions from wild teosinte to modern maize, with resistant varieties maintaining key bacterial metabolic pathways that may contribute to WCR resistance.

## DISCUSSION

Our findings showed that resistance to WCR in teosinte and maize involves specialized interactions between plant defense mechanisms and rhizosphere microbiome dynamics, with particularly robust responses observed in teosinte. Through a comprehensive analysis of WCR and plant performance metrics and rhizosphere community dynamics, we identified complex defense networks that integrate direct plant responses and microbiome-mediated protection. The superior resistance observed in wild teosinte and ancestral maize accessions, exemplified by Los Naranjos teosinte (overall performance score: 0.8915) and Tehua ancestral maize (0.8461), suggests the preservation of ancestral resistance mechanisms that may have been silenced with breeding [8,14].

The quantitative assessment of larval development patterns provides compelling evidence for resistance mechanisms operating in wild teosinte. Resistant wild teosinte accessions effectively impeded larval progression, with 70–90% of larvae remaining in first and second instars (≤0.06 cm head capsule width), comparable to resistance levels reported in other wild crop relatives (73–82% early instar retention) [59,60]. This developmental arrest was coupled with significant reductions in average larval weight (1.8 ± 0.4 mg in resistant vs. 7.2 ± 1.2 mg in susceptible accessions; *p* < 0.001) and markedly lower survival rates (22.5% ± 4.8% in resistant vs. 78.6% ± 6.3% in susceptible genotypes; *p* < 0.001) (Fig. S1), indicating strong antibiosis effects. These results strongly support the hypothesis that resistant teosinte lines deploy multiple, coordinated defense barriers that interfere with herbivore development and survival.

Our canonical centroid analysis, which explained 86.15% of total variance (PC1: 64.74%, PC2: 21.41%), clearly separated resistance categories and highlighted contrasting strategies between resistant, intermediately resistant, and susceptible groups. Significant differences in biomass reduction (*p* = 0.0028), larval biomass (*p* = 0.0226), and survival further underscored these distinctions. Intermediately resistant genotypes displayed intermediate levels of infestation (*p* = 0.0038) and biomass loss, suggesting that partial resistance may involve compensatory mechanisms or reduced attractiveness to larvae [64,65]. These patterns align with previous findings in other wild relatives, such as *Solanum* spp., where herbivore resistance is often maintained through conserved traits such as trichome density, defensive metabolites, and microbe-mediated antibiosis [62,63,66,67]. The retention of such characteristics in teosinte offers a valuable model for reintroducing lost resistance components into modern maize cultivars.

Perhaps our most striking finding was the correlation between plant resistance and rhizosphere microbiome composition. Alpha diversity analyses revealed significantly higher microbial diversity in resistant accessions under WCR herbivory (*p* < 0.001), with accessions like the resistant wild teosinte (Los Naranjos) exhibiting substantial increases in OTU richness, phylogenetic diversity, and Shannon diversity following WCR infestation. Specifically, this teosinte showed marked increases in OTUs (*p* < 0.0001), phylogenetic diversity (*p* < 0.0001), and Shannon index (*p* < 0.0001). In contrast, the susceptible wild teosinte (Guachinango) showed limited or non-significant responses (*p* = 0.003, 0.012, and ns, respectively). This enhanced diversity suggests a more complex and resilient microbial network among teosintes that may bolster host defense under herbivory [39, 68]. Overall, the strong community differentiation observed in the UniFrac analysis (PC1: 84.61%) between WCR-infested and control plants indicated substantial restructuring of microbial communities in response to WCR herbivory, with community shifts occurring consistently across all plant groups, regardless of resistance level.

The co-enrichment patterns observed in rhizosphere bacterial communities provide compelling evidence for structured microbial consortia development in resistant genotypes. The resistant wild teosinte (Los Naranjos) harbored 47 bacterial species, with 31 significantly enriched following WCR herbivory, compared to only 7 enriched species in the susceptible teosinte (Guachinango). Notable enriched taxa in resistant genotypes included *Pseudomonas putida* (3.0-fold), *Stenotrophomonas maltophilia* (2.8-fold), and *Enterobacter tabaci* (2.5-fold), supporting the hypothesis of active recruitment of beneficial Proteobacteria [70, 71]. These enrichment patterns mirror those documented in disease-suppressive soils, where similar taxonomic groups exhibit consistent abundance shifts in response to biotic pressure [72, 73, 74].

Analysis of predicted microbial functions revealed sophisticated defense-related metabolic networks in resistant teosinte. Functional pathways such as antibiotic production (log FC = 3.2), hydrogen cyanide production (log FC = 2.6), DAPG synthesis (log FC = 2.8), benzoxazinoid metabolism (log FC = 2.9), and chitinase production (log FC = 2.4) were significantly enriched, indicating a coordinated microbiome response to insect herbivory [75, 76]. The maintenance of metabolic network integrity under WCR herbivory in resistant accessions (contrasting with the functional disruption seen in susceptible accessions) suggests the presence of evolved mechanisms for microbial community stability and resilience [77, 78]. Resistant accessions also maintained functions such as quorum sensing regulation (log FC = 3.5), protease inhibitors (log FC = 2.5), and pyoverdine synthesis (log FC = 2.7), reinforcing the role of microbial regulation in defense [79, 80].

The remarkable performance of teosinte accessions, particularly Los Naranjos (our resistant wild teosinte), reflects the preservation of highly regulated, multi-layered defense strategies that integrate plant-intrinsic resistance with microbiome-mediated protection. Los Naranjos teosinte exhibited superior resistance metrics (including reduced larval development, enhanced microbial diversity, and the highest number of enriched defensive taxa), highlighting the potential of ancestral traits for modern applications [81, 82]. This contrasts sharply with modern cultivars like B73, which showed fewer enriched functions and lower microbial responsiveness, suggesting that domestication and breeding have significantly eroded these complex resistance traits [83, 84, 85].

These findings have important implications for the development of microbial products targeting WCR and other pests, as well as breeding strategies to enhance WCR resistance in maize. On one hand, several microbial taxa consistently enriched in resistant accessions, such as *Pseudomonas putida* (3.0-3.2-fold), *Stenotrophomonas maltophilia* (2.7-2.9-fold), and *Bacillus subtilis*, have well-documented insecticidal properties and could be developed as biocontrol inoculants to manage WCR. On the other hand, “microbiome-smart” breeding programs could enhance pest resistance in maize by integrating selection for plants that recruit, promote, and sustain microbial consortia relevant to WCR management [86, 87]. The consistent ability of resistant-group accessions to enrich 28–31 beneficial bacterial species (especially within Proteobacteria) compared to 7–19 beneficial species in susceptible-group accessions offers a viable target for trait selection [88, 89, 90]. Incorporating microbiome-mediated resistance into modern maize varieties offers multiple sustainability benefits. In this study, resistant accessions maintained higher microbial diversity, stability under stress, and functional enrichment hallmarks of resilient systems. These qualities enhanced WCR resistance and may improve soil health and long-term ecosystem functioning [91, 92]. The wild teosinte accessions evidenced how maintaining microbial diversity (e.g., OTU increases of >40 units under WCR herbivory) and network integrity can lead to stable plant-microbiome alliances even under biotic stress [93, 94].

Our results suggest several promising research directions. Firstly, unraveling the genetic architecture underlying the differential recruitment of beneficial microbiota may yield novel resistance loci or root exudate pathways for breeding [95, 96]. The consistent upregulation of key defense-related functions (log FC range: 2.4–3.5) in resistant accessions points to robust regulatory circuits worthy of transcriptomic and metabolomic dissection [97, 98]. Additionally, synthetic microbiome engineering based on the enriched taxa in resistant lines, coupled with agronomic practices to favor their establishment, offers a promising path forward [99, 100]. Altogether, our results add to existing research that points to the promise of shifting toward integrated pest management strategies that recognize the synergistic potential between plant genotypic traits and microbiota, and how these interactions mediate plant survival and reproduction, i.e., productivity in agriculture [101, 102]. Further research examining interactions between crop breeding lines and beneficial microbiota could revolutionize crop breeding and pest control and lead to greater resilience and sustainability in agriculture [103, 104]. As the demand grows for climate-smart and environmentally friendly agricultural technologies, it seems that exploiting the evolutionarily shaped solutions for overcoming stress observed in teosinte-microbiome alliances could serve as a blueprint for future innovations in crop protection [105, 106].

## CONCLUSION

Our findings reveal that the rhizosphere microbiome plays a critical role in shaping resistance responses to *Diabrotica virgifera virgifera* herbivory in teosinte, and to some extent in maize. Resistant genotypes consistently exhibited increased microbial diversity, and under WCR herbivory, displayed significant restructuring of rhizosphere communities and selective enrichment of functional taxa and metabolic pathways associated with plant defense against herbivores. These results highlight a coordinated defense system integrating plant genetic traits and microbiome-mediated mechanisms. These changes were absent or attenuated in susceptible accessions of wild teosinte, ancestral maize, and modern maize, indicating that resistance is associated not only with the recruitment of beneficial taxa, such as *Pseudomonas* and *Stenotrophomonas*, but also with the maintenance of functional network stability under biotic stress. The preservation of such traits in wild teosinte, particularly in the resistant accession (Los Naranjos), underscores the potential of leveraging ancestral microbiome interactions for crop improvement. Incorporating microbiome-informed approaches into breeding programs may enable the development of maize cultivars with enhanced and durable resistance to WCR, contributing to sustainable pest management and agroecosystem resilience.

## ACKNOWLEDGMENTS

We thank undergraduate students Arturo España and Amrita Gabu, as well as graduate students Nihad Kerrour, John Grunseich, and Ilksen Topcu for providing technical support in the laboratory.

## AUTHOR CONTRIBUTIONS

Conceptualization: J.S.B., E.D.V.C., S.A.B., Data curation: E.D.V.C, C.H.H.R., S.A.B., Formal analysis: E.D.V.C. Funding acquisition: J.S.B., S.A.B., Investigation: E.D.V.C., C.H.H.R., J.S.B., S.A.B. Methodology: S.A.B., J.S.B., Project administration: S.A.B., J.S.B., Resources: S.A.B., J.S.B., Software: E.D.V.C. Supervision: S.A.B., J.S.B., Validation: S.A.B., J.S.B., Visualization: E.D.V.C., Writing – original draft: E.D.V.C. Writing – review & editing: E.D.V.C., C.H.H.R., J.S.B., S.A.B.

## REFERENCES

1. De Lange, E. S., Balmer, D., Mauch Mani, B., and Turlings, T. C. 2014. Insect and pathogen attack and resistance in maize and its wild ancestors, the teosintes. New Phytol. 204:329–341. doi:10.1111/nph.13005

2. Meinke, L. J., Sappington, T. W., Onstad, D. W., Guillemaud, T., Miller, N. J., Komáromi, J., Levay, N., Kiss, J., Toth, F., Toth, M., Toepfer, S., and Kuhlmann, U. 2009. Western corn rootworm (Diabrotica virgifera virgifera LeConte) population dynamics. Agric. For. Entomol. 11:29–46. doi:10.1111/j.1461-9563.2008.00419.x

3. Gray, M. E., Sappington, T. W., Miller, N. J., Moeser, J., and Bohn, M. O. 2009. Adaptation and invasiveness of western corn rootworm: intensifying research on a worsening pest. Annu. Rev. Entomol. 54:303–321. doi:10.1146/annurev.ento.54.110807.090434

4. Romeis, J., Meissle, M., and Bigler, F. 2006. Transgenic crops expressing Bacillus thuringiensis toxins and biological control. Nat. Biotechnol. 24:63–71. doi:10.1038/nbt1180

5. Gassmann, A. J., Petzold-Maxwell, J. L., Keweshan, R. S., and Dunbar, M. W. 2011. Field-evolved resistance to Bt maize by western corn rootworm. PLoS ONE 6:e22629. doi:10.1371/journal.pone.0022629

6. Jurat-Fuentes, J. L., Heckel, D. G., and Ferré, J. 2021. Mechanisms of resistance to insecticidal proteins from Bacillus thuringiensis. Annu. Rev. Entomol. 66:121–140. doi:10.1146/annurev-ento-052620-073348

7. Maag, D., Erb, M., Köllner, T. G., and Gershenzon, J. 2015. Defensive weapons and defense signals in plants: Some metabolites serve both roles. BioEssays 37:167–174. doi:10.1002/bies.201400124

8. Fontes-Puebla, A. A., and Bernal, J. S. 2020. Resistance and tolerance to root herbivory in maize were mediated by domestication, spread, and breeding. Front. Plant Sci. 11:223. doi:10.3389/fpls.2020.00223

9. Erb, M., Glauser, G., and Robert, C. A. 2012. Induced immunity against belowground insect herbivores-activation of defenses in the absence of a jasmonate burst. J. Chem. Ecol. 38:629–640. doi:10.1007/s10886-012-0107-9

10. Robert, C. A. M., Erb, M., Duployer, M., Zwahlen, C., Doyen, G. R., and Turlings, T. C. J. 2012. Herbivore-induced plant volatiles mediate host selection by a root herbivore. New Phytol. 194:1061–1069. doi:10.1111/j.1469-8137.2012.04127.x

11. Trivedi, P., Leach, J. E., Tringe, S. G., Sa, T., and Singh, B. K. 2020. Plant–microbiome interactions: from community assembly to plant health. Nat. Rev. Microbiol. 18:607–621. doi:10.1038/s41579-020-0412-1

12. Rodriguez, P. A., Rothballer, M., Chowdhury, S. P., Nussbaumer, T., Gutjahr, C., and Falter-Braun, P. 2019. Systems biology of plant-microbiome interactions. Mol. Plant 12:804–821. doi:10.1016/j.molp.2019.05.006

13. Berendsen, R. L., Pieterse, C. M., and Bakker, P. A. 2012. The rhizosphere microbiome and plant health. Trends Plant Sci. 17:478–486. doi:10.1016/j.tplants.2012.04.001

14. Hu, L., Robert, C. A. M., Cadot, S., Zhang, X., Ye, M., Li, B., Manzo, D., Chervet, N., Steinger, T., van der Heijden, M. G. A., Schlaeppi, K., and Erb, M. 2018. Root exudate metabolites drive plant-soil feedback on growth and defense by shaping the rhizosphere microbiota. Nat. Commun. 9:2738. doi:10.1038/s41467-018-05122-7

15. Erb, M., Flors, V., Karlen, D., De Lange, E., Planchamp, C., D’Alessandro, M., Turlings, T. C. J., and Ton, J. 2009. Signal signature of aboveground-induced resistance upon belowground herbivory in maize. Plant J. 59:292–302. doi:10.1111/j.1365-313X.2009.03868.x

16. Carrière, Y., Crickmore, N., and Tabashnik, B. E. 2015. Optimizing pyramided transgenic Bt crops for sustainable pest management. Nat. Biotechnol. 33:161–168. doi:10.1038/nbt.3099

17. Meihls, L. N., Higdon, M. L., Siegfried, B. D., Miller, N. J., Sappington, T. W., Ellersieck, M. R., Spencer, T. A., and Hibbard, B. E. 2008. Increased survival of western corn rootworm on transgenic corn within three generations of on-plant greenhouse selection. Proc. Natl. Acad. Sci. USA 105:19177–19182. doi:10.1073/pnas.0805565105

18. Martínez-Medina, A., Flors, V., Heil, M., Mauch-Mani, B., Pieterse, C. M., Pozo, M. J., Ton, J., van Dam, N. M., and Conrath, U. 2016. Recognizing plant defense priming. Trends Plant Sci. 21:818–822. doi:10.1016/j.tplants.2016.07.009

19. Pérez-Jaramillo, J. E., Mendes, R., and Raaijmakers, J. M. 2016. Impact of plant domestication on rhizosphere microbiome assembly and functions. Plant Mol. Biol. 90:635–644. doi:10.1007/s11103-015-0337-7

20. Köllner, T. G., Held, M., Lenk, C., Hiltpold, I., Turlings, T. C., Gershenzon, J., and Degenhardt, J. 2008. A maize (E)-β-caryophyllene synthase implicated in indirect defense responses against herbivores is not expressed in most American maize varieties. Plant Cell 20:482–494. doi:10.1105/tpc.107.051672

21. Rasmann, S., Köllner, T. G., Degenhardt, J., Hiltpold, I., Toepfer, S., Kuhlmann, U., Gershenzon, J., and Turlings, T. C. J. 2005. Recruitment of entomopathogenic nematodes by insect-damaged maize roots. Nature 434:732–737. doi:10.1038/nature03451

22. Bulgarelli, D., Schlaeppi, K., Spaepen, S., Van Themaat, E. V. L., and Schulze-Lefert, P. 2013. Structure and functions of the bacterial microbiota of plants. Annu. Rev. Plant Biol. 64:807–838. doi:10.1146/annurev-arplant-050312-120106

23. Lundgren, J. G., and Fausti, S. W. 2015. Trading biodiversity for pest problems. Sci. Adv. 1:e1500558. doi:10.1126/sciadv.1500558

24. Marti, G., Erb, M., Boccard, J., Glauser, G., Doyen, G. R., Villard, N., Robert, C. A. M., Turlings, T. C. J., Wolfender, J. L., and Rudaz, S. 2013. Metabolomics reveals herbivore-induced metabolites of resistance and susceptibility in maize leaves and roots. Plant Cell Environ. 36:621–639. doi:10.1111/pce.12002

25. Christensen, S. A., Nemchenko, A., Borrego, E., Murray, I., Sobhy, I. S., Bosak, L., DeBlasio, S., Erb, M., Robert, C. A. M., Vaughn, K. C., Herrfurth, C., Tumlinson, J., Feussner, I., Jackson, D., Turlings, T. C. J., Engelberth, J., Nansen, C., Meeley, R., and Kolomiets, M. V. 2013. The maize lipoxygenase, ZmLOX10, mediates green leaf volatile, jasmonate and herbivore induced plant volatile production for defense against insect attack. Plant J. 74:59–73. doi:10.1111/tpj.12101

26. Chinchilla, D., Bruisson, S., Meyer, S., Zühlke, D., Riedel, K., and Weisskopf, L. 2019. A sulfur-containing volatile emitted by potato-associated bacteria confers protection against late blight through direct anti-oomycete activity. Sci. Rep. 9:18778. doi:10.1038/s41598-019-55218-3

27. Gaillard, M. D. P., Glauser, G., Robert, C. A. M., and Turlings, T. C. J. 2018. Fine-tuning the ‘plant domestication-reduced defense’ hypothesis: specialist vs generalist herbivores. New Phytol. 217:355–366. doi:10.1111/nph.14757

28. Aguirre-Liguori, J. A., Ramírez-Barahona, S., Tiffin, P., and Eguiarte, L. E. 2019. Climate change is predicted to disrupt patterns of local adaptation in wild and cultivated maize. Proc. R. Soc. B 286:20190486. doi:10.1098/rspb.2019.0486

29. Szczepaniec, A., Widney, S. E., Bernal, J. S., and Eubanks, M. D. 2013. Higher expression of induced defenses in teosintes (Zea spp.) is correlated with greater resistance to fall armyworm Spodoptera frugiperda Smith (Lepidoptera: Noctuidae). Entomol. Exp. Appl. 146:242–251. doi:10.1111/eea.12014

30. Grunseich, J. M., Huang, P. C., Bernal, J. S., and Kolomiets, M. 2025. Western corn rootworm resistance in maize persists in the absence of jasmonic acid. Planta 261:6. doi:10.1007/s00425-024-04580-2

31. Lundberg, D. S., Lebeis, S. L., Paredes, S. H., Yourstone, S., Gehring, J., Malfatti, S., Tremblay, J., Engelbrektson, A., Kunin, V., del Rio, T. G., Edgar, R. C., Eickhorst, T., Ley, R. E., Hugenholtz, P., Tringe, S. G., and Dangl, J. L. 2012. Defining the core Arabidopsis thaliana root microbiome. Nature 488:86–90. doi:10.1038/nature11237

32. Hufford, M. B., Bilinski, P., Pyhäjärvi, T., and Ross-Ibarra, J. 2012. Teosinte as a model system for population and ecological genomics. Trends Genet. 28:606–615. doi:10.1016/j.tig.2012.08.004

33. Robert, C. A. M., Erb, M., Hibbard, B. E., French, B. W., Zwahlen, C., and Turlings, T. C. J. 2012. A specialist root herbivore reduces plant resistance and uses an induced plant volatile to aggregate in a density dependent manner. Funct. Ecol. 26:1429–1440. doi:10.1111/j.1365-2435.2012.02030.x

34. Schumann, M., Patel, A., and Vidal, S. 2014. Soil application of an encapsulated CO2 source and its potential for management of western corn rootworm larvae. J. Econ. Entomol. 107:230–239. doi:10.1603/EC13344

35. Erb, M., Robert, C. A. M., Hibbard, B. E., and Turlings, T. C. J. 2011. Sequence of arrival determines plant mediated interactions between herbivores. J. Ecol. 99:7–15. doi:10.1111/j.1365-2745.2010.01757.x

36. Robert, C. A. M., Veyrat, N., Glauser, G., Marti, G., Doyen, G. R., Villard, N., Gaillard, M. D. P., Köllner, T. G., Giron, D., Body, M., Babst, B. A., Ferrieri, R. A., Turlings, T. C. J., and Erb, M. 2012. A specialist root herbivore exploits defensive metabolites to locate nutritious tissues. Ecol. Lett. 15:55–64. doi:10.1111/j.1461-0248.2011.01708.x

37. Edwards, J. E., Johnson, C., Santos-Medellín, C., Lurie, E., Podishetty, N. K., Bhatnagar, S., Eisen, J. A., and Sundaresan, V. 2015. Structure, variation, and assembly of the root-associated microbiomes of rice. Proc. Natl. Acad. Sci. USA 112:E911–E920. doi:10.1073/pnas.1414592112

38. Lundberg, D. S., Yourstone, S., Mieczkowski, P., Jones, C. D., and Dangl, J. L. 2013. Practical innovations for high-throughput amplicon sequencing. Nat. Methods 10:999–1002. doi:10.1038/nmeth.2634

39. Lebeis, S. L., Paredes, S. H., Lundberg, D. S., Breakfield, N., Gehring, J., McDonald, M., Malfatti, S., del Rio, T. G., Jones, C. D., Tringe, S. G., and Dangl, J. L. 2015. Salicylic acid modulates colonization of the root microbiome by specific bacterial taxa. Science 349:860–864. doi:10.1126/science.aaa8764

40. Gassmann, A. J., Petzold-Maxwell, J. L., Clifton, E. H., Dunbar, M. W., Hoffmann, A. M., Ingber, D. A., and Keweshan, R. S. 2014. Field-evolved resistance by western corn rootworm to multiple Bacillus thuringiensis toxins in transgenic maize. Proc. Natl. Acad. Sci. USA 111:5141–5146. doi:10.1073/pnas.1317179111

41. Hiltpold, I., Hibbard, B. E., French, B. W., and Turlings, T. C. J. 2012. Capsules containing entomopathogenic nematodes as a Trojan horse approach to control the western corn rootworm. Plant Soil 358:11–25. doi:10.1007/s11104-012-1253-0

42. Robert, C. A. M., Schirmer, S., Barry, J., French, B. W., Hibbard, B. E., and Gershenzon, J. 2015. Belowground herbivore tolerance involves delayed root growth and resource allocation. Plant Cell 27:2350–2363. doi:10.1105/tpc.15.00006

43. Machado, R. A. R., Robert, C. A. M., Arce, C. C. M., Ferrieri, A. P., Xu, S., Jimenez-Aleman, G. H., Baldwin, I. T., and Erb, M. 2016. Auxin is rapidly induced by herbivore attack and regulates a subset of systemic, jasmonate-dependent defenses. Plant Physiol. 172:521–532. doi:10.1104/pp.16.00940

44. Toepfer, S., Kuhlmann, U., and Jehle, J. A. 2006. Diagnostic tools to determine virulence of Diabrotica-effective entomopathogenic nematodes. IOBC/WPRS Bull. 29:159–165.

45. Kurtz, B., Karlovsky, P., and Vidal, S. 2010. Interaction between western corn rootworm (Coleoptera: Chrysomelidae) larvae and root-infecting Fusarium verticillioides. Environ. Entomol. 39:1532–1538. doi:10.1603/EN10025

46. Hammack, L., Ellsbury, M. M., Roehrdanz, R. L., and Pikul, J. L. 2003. Larval sampling and instar determination in field populations of northern and western corn rootworm (Coleoptera: Chrysomelidae). J. Econ. Entomol. 96:1153–1159. doi:10.1093/jee/96.4.1153

47. Zhalnina, K., Louie, K. B., Hao, Z., Mansoori, N., da Rocha, U. N., Shi, S., Cho, H., Karaoz, U., Loqué, D., Bowen, B. P., Firestone, M. K., Northen, T. R., and Brodie, E. L. 2018. Dynamic root exudate chemistry and microbial substrate preferences drive patterns in rhizosphere microbial community assembly. Nat. Microbiol. 3:470–480. doi:10.1038/s41564-018-0129-3

48. Walters, W. A., Jin, Z., Youngblut, N., Wallace, J. G., Sutter, J., Zhang, W., González-Peña, A., Peiffer, J., Koren, O., Shi, Q., Knight, R., del Rio, T. G., Tringe, S. G., Buckler, E. S., Dangl, J. L., and Ley, R. E. 2018. Large-scale replicated field study of maize rhizosphere identifies heritable microbes. Proc. Natl. Acad. Sci. USA 115:7368–7373. doi:10.1073/pnas.1800918115

49. Fiedorová, K., Radvanský, M., Němcová, E., Grombiříková, H., Bosák, J., Černochová, M., Lexa, M., Šmajs, D., and Freiberger, T. 2019. The impact of DNA extraction methods on stool bacterial and fungal microbiota community recovery. Front. Microbiol. 10:821. doi:10.3389/fmicb.2019.00821

50. Wick, R. R., Judd, L. M., and Holt, K. E. 2019. Performance of neural network basecalling tools for Oxford Nanopore sequencing. Genome Biol. 20:129. doi:10.1186/s13059-019-1727-y

51. De Coster, W., D’Hert, S., Schultz, D. T., Cruts, M., and Van Broeckhoven, C. 2018. NanoPack: visualizing and processing long-read sequencing data. Bioinformatics 34:2666–2669. doi:10.1093/bioinformatics/bty149

52. Chen, S., Zhou, Y., Chen, Y., and Gu, J. 2018. fastp: an ultra-fast all-in-one FASTQ preprocessor. Bioinformatics 34:884–890. doi:10.1093/bioinformatics/bty560

53. Li, H. 2018. Minimap2: pairwise alignment for nucleotide sequences. Bioinformatics 34:3094–3100. doi:10.1093/bioinformatics/bty191

54. Anderson, M. J. 2017. Permutational Multivariate Analysis of Variance (PERMANOVA). Wiley StatsRef: Statistics Reference Online 1–15. doi:10.1002/9781118445112.stat07841

55. McMurdie, P. J., and Holmes, S. 2013. phyloseq: An R Package for Reproducible Interactive Analysis and Graphics of Microbiome Census Data. PLoS ONE 8:e61217. doi:10.1371/journal.pone.0061217

56. Louca, S., Parfrey, L. W., and Doebeli, M. 2016. Decoupling function and taxonomy in the global ocean microbiome. Science 353:1272–1277. doi:10.1126/science.aaf4507

57. Douglas, G. M., Maffei, V. J., Zaneveld, J. R., Yurgel, S. N., Brown, J. R., Taylor, C. M., Huttenhower, C., and Langille, M. G. I. 2020. PICRUSt2 for prediction of metagenome functions. Nat. Biotechnol. 38:685–688. doi:10.1038/s41587-020-0548-6

58. Benjamini, Y., and Hochberg, Y. 1995. Controlling the False Discovery Rate: A Practical and Powerful Approach to Multiple Testing. J. R. Stat. Soc. Series B Stat. Methodol. 57:289–300. doi:10.1111/j.2517-6161.1995.tb02031.x

59. Xu, L., Naylor, D., Dong, Z., Simmons, T., Pierroz, G., Hixson, K. K., Kim, Y. M., Zink, E. M., Engbrecht, K. M., Wang, Y., Gao, C., DeGraaf, S., Madera, M. A., Sievert, J. A., Hollingsworth, J., Birdseye, D., Scheller, H. V., Hutmacher, R., Dahlberg, J., Jansson, C., Taylor, J. W., Vogel, J. P., Lemaux, P. G., Clay, K., Hutmacher, R. B., Coaldrake, P., Dahlberg, J., Lemaux, P. G., and Coleman-Derr, D. 2018. Drought delays development of the sorghum root microbiome and enriches for monoderm bacteria. Proc. Natl. Acad. Sci. USA 115:E4284–E4293. doi:10.1073/pnas.1717308115

60. Moreira, X., Abdala-Roberts, L., Gols, R., and Francisco, M. 2018. Plant domestication decreases both constitutive and induced chemical defences by direct selection against defensive traits. Sci. Rep. 8:12678. doi:10.1038/s41598-018-31041-0

61. Zhu, S., Vivanco, J. M., and Manter, D. K. 2016. Nitrogen fertilizer rate affects root exudation, the rhizosphere microbiome and nitrogen-use-efficiency of maize. Appl. Soil Ecol. 107:324–333. doi:10.1016/j.apsoil.2016.07.009

62. Schuman, M. C., and Baldwin, I. T. 2016. The layers of plant responses to insect herbivores. Annu. Rev. Entomol. 61:373–394. doi:10.1146/annurev-ento-010715-023851

63. Turlings, T. C. J., and Erb, M. 2018. Tritrophic interactions mediated by herbivore-induced plant volatiles: mechanisms, ecological relevance, and application potential. Annu. Rev. Entomol. 63:433–452. doi:10.1146/annurev-ento-020117-043507

64. Kessler, A., and Heil, M. 2011. The multiple faces of indirect defences and their agents of natural selection. Funct. Ecol. 25:348–357. doi:10.1111/j.1365-2435.2010.01818.x

65. Salgado Luarte, C., González Teuber, M., Madriaza, K., and Gianoli, E. 2023. Trade off between plant resistance and tolerance to herbivory: mechanical defenses outweigh chemical defenses. Ecology 104:e3860. doi:10.1002/ecy.3860

66. Chen, Y. H., Gols, R., and Benrey, B. 2015. Crop domestication and its impact on naturally selected trophic interactions. Annu. Rev. Entomol. 60:35–58. doi:10.1146/annurev-ento-010814-020601

67. Walling, L. L. 2008. Avoiding effective defenses: strategies employed by phloem-feeding insects. Plant Physiol. 146:859–866. doi:10.1104/pp.107.113142

68. Tkacz, A., Cheema, J., Chandra, G., Grant, A., and Poole, P. S. 2015. Stability and succession of the rhizosphere microbiota depends upon plant type and soil composition. ISME J. 9:2349–2359. doi:10.1038/ismej.2015.41

69. Mendes, R., Garbeva, P., and Raaijmakers, J. M. 2013. The rhizosphere microbiome: significance of plant beneficial, plant pathogenic, and human pathogenic microorganisms. FEMS Microbiol. Rev. 37:634–663. doi:10.1111/1574-6976.12028

70. Mendes, R., Kruijt, M., De Bruijn, I., Dekkers, E., Van Der Voort, M., Schneider, J. H. M., Piceno, Y. M., DeSantis, T. Z., Andersen, G. L., Bakker, P. A. H. M., and Raaijmakers, J. M. 2011. Deciphering the rhizosphere microbiome for disease-suppressive bacteria. Science 332:1097–1100. doi:10.1126/science.1203980

71. Pascale, A., Proietti, S., Pantelides, I. S., and Stringlis, I. A. 2020. Modulation of the root microbiome by plant molecules: the basis for targeted disease suppression and plant growth promotion. Front. Plant Sci. 10:1741. doi:10.3389/fpls.2019.01741

72. Carrión, V. J., Perez-Jaramillo, J., Cordovez, V., Tracanna, V., De Hollander, M., Ruiz-Buck, D., Mendes, L. W., van Ijcken, W. F. J., Gomez-Exposito, R., Elsayed, S. S., Mohanraju, P., Arifah, A., van der Ent, S., van der Hooft, J. J. J., Alonso-Blanco, C., Bai, Y., Bakker, P. A. H. M., Oome, S., Cornellissen, B. J. C., Raaijmakers, J. M., and Medema, M. H. 2019. Pathogen-induced activation of disease-suppressive functions in the endophytic root microbiome. Science 366:606–612. doi:10.1126/science.aaw9285

73. Durán, P., Thiergart, T., Garrido-Oter, R., Agler, M., Kemen, E., Schulze-Lefert, P., and Hacquard, S. 2018. Microbial interkingdom interactions in roots promote Arabidopsis survival. Cell 175:973–983. doi:10.1016/j.cell.2018.10.020

74. de Boer, W. 2017. Upscaling of fungal–bacterial interactions: from the lab to the field. Curr. Opin. Microbiol. 37:35–41. doi:10.1016/j.mib.2017.03.007

75. Peiffer, J. A., Spor, A., Koren, O., Jin, Z., Tringe, S. G., Dangl, J. L., Buckler, E. S., and Ley, R. E. 2013. Diversity and heritability of the maize rhizosphere microbiome under field conditions. Proc. Natl. Acad. Sci. USA 110:6548–6553. doi:10.1073/pnas.1302837110

76. Pineda, A., Kaplan, I., and Bezemer, T. M. 2017. Steering soil microbiomes to suppress aboveground insect pests. Trends Plant Sci. 22:770–778. doi:10.1016/j.tplants.2017.07.002

77. Lugtenberg, B., and Kamilova, F. 2009. Plant-growth-promoting rhizobacteria. Annu. Rev. Microbiol. 63:541–556. doi:10.1146/annurev.micro.62.081307.162918

78. Bai, Y., Müller, D. B., Srinivas, G., Garrido-Oter, R., Potthoff, E., Rott, M., Dombrowski, N., Münch, P. C., Spaepen, S., Remus-Emsermann, M., Hüttel, B., McHardy, A. C., Vorholt, J. A., and Schulze-Lefert, P. 2015. Functional overlap of the Arabidopsis leaf and root microbiota. Nature 528:364–369. doi:10.1038/nature16192

79. Stringlis, I. A., Yu, K., Feussner, K., de Jonge, R., Van Bentum, S., Van Verk, M. C., Berendsen, R. L., Bakker, P. A. H. M., Bloem, E., Dijkstra, P., Fukushima, A., Kalkhoven, M. A., Kistenkas, M., Poelman, E. H., Raaijmakers, J. M., Takken, F. L. W., Wees, S. C. M., Van Wees, S. C. M., Vlot, A. C., Wu, X., Pieterse, C. M. J., and Van Pelt, J. A. 2018. MYB72-dependent coumarin exudation shapes root microbiome assembly to promote plant health. Proc. Natl. Acad. Sci. USA 115:E5213–E5222. doi:10.1073/pnas.1722335115

80. Cotton, T. E. A., Pétriacq, P., Cameron, D. D., Meselmani, M. A., Schwarzenbacher, R., Rolfe, S. A., and Ton, J. 2019. Metabolic regulation of the maize rhizobiome by benzoxazinoids. ISME J. 13:1647–1658. doi:10.1038/s41396-019-0375-2

81. Hufford, M. B., Xu, X., van Heerwaarden, J., Pyhäjärvi, T., Chia, J. M., Cartwright, R. A., Elshire, R. J., Glaubitz, J. C., Guill, K. E., Kaeppler, S. M., Lai, J., Morrell, P. L., Shannon, L. M., Song, C., Springer, N. M., Swanson-Wagner, R. A., Tiffin, P., Wang, J., Zhang, G., Doebley, J., McMullen, M. D., Ware, D., Buckler, E. S., Yang, S., and Ross-Ibarra, J. 2012. Comparative population genomics of maize domestication and improvement. Nat. Genet. 44:808–811. doi:10.1038/ng.2309

82. Chen, Y. H., Gols, R., Stratton, C. A., Brevik, K. A., and Benrey, B. 2020. Complex evolutionary history of the Mexican wild bean Phaseolus vulgaris reveals parallel signatures of domestication in mesoamerican and andean populations. Front. Plant Sci. 11:571352. doi:10.3389/fpls.2020.571352

83. Martínez-Medina, A., Fernández, I., Sánchez-Guzmán, M. J., Jung, S. C., Pascual, J. A., and Pozo, M. J. 2013. Deciphering the hormonal signalling network behind the systemic resistance induced by Trichoderma harzianum in tomato. Front. Plant Sci. 4:206. doi:10.3389/fpls.2013.00206

84. Wei, Z., and Jousset, A. 2017. Plant breeding goes microbial. Trends Plant Sci. 22:555–558. doi:10.1016/j.tplants.2017.05.009

85. Cordovez, V., Dini-Andreote, F., Carrión, V. J., and Raaijmakers, J. M. 2019. Ecology and evolution of plant microbiomes. Annu. Rev. Microbiol. 73:69–88. doi:10.1146/annurev-micro-090817-062524

86. Sanchez-Canizares, C., Jorrín, B., Poole, P. S., and Tkacz, A. 2017. Understanding the holobiont: the interdependence of plants and their microbiome. Curr. Opin. Microbiol. 38:188–196. doi:10.1016/j.mib.2017.07.001

87. Wille, L., Messmer, M. M., Studer, B., and Hohmann, P. 2019. Insights to plant–microbe interactions provide opportunities to improve resistance breeding against root diseases in grain legumes. Plant Cell Environ. 42:20–40. doi:10.1111/pce.13214

88. Mueller, U. G., and Sachs, J. L. 2015. Engineering microbiomes to improve plant and animal health. Trends Microbiol. 23:606–617. doi:10.1016/j.tim.2015.07.009

89. Toju, H., Peay, K. G., Yamamichi, M., Narisawa, K., Hiruma, K., Naito, K., Fukuda, S., Ushio, M., Nakaoka, S., Onoda, Y., Yoshida, K., Schlaeppi, K., Bai, Y., Sugiura, R., Ichihashi, Y., Minamisawa, K., and Kiers, E. T. 2018. Core microbiomes for sustainable agroecosystems. Nat. Plants 4:247–257. doi:10.1038/s41477-018-0139-4

90. Oyserman, B. O., Medema, M. H., and Raaijmakers, J. M. 2018. Road MAPs to engineer host microbiomes. Curr. Opin. Microbiol. 43:46–54. doi:10.1016/j.mib.2017.11.023

91. Friesen, M. L., Porter, S. S., Stark, S. C., von Wettberg, E. J., Sachs, J. L., and Martinez-Romero, E. 2011. Microbially mediated plant functional traits. Annu. Rev. Ecol. Evol. Syst. 42:23–46. doi:10.1146/annurev-ecolsys-102710-145039

92. Busby, P. E., Soman, C., Wagner, M. R., Friesen, M. L., Kremer, J., Bennett, A., Morsy, M., Eisen, J. A., Leach, J. E., and Dangl, J. L. 2017. Research priorities for harnessing plant microbiomes in sustainable agriculture. PLoS Biol. 15:e2001793. doi:10.1371/journal.pbio.2001793

93. Chaparro, J. M., Sheflin, A. M., Manter, D. K., and Vivanco, J. M. 2012. Manipulating the soil microbiome to increase soil health and plant fertility. Biol. Fertil. Soils 48:489–499. doi:10.1007/s00374-012-0691-4

94. Finkel, O. M., Castrillo, G., Herrera Paredes, S., Salas González, I., and Dangl, J. L. 2017. Understanding and exploiting plant beneficial microbes. Curr. Opin. Plant Biol. 38:155–163. doi:10.1016/j.pbi.2017.04.018

95. Wagner, M. R., Lundberg, D. S., Coleman-Derr, D., Tringe, S. G., Dangl, J. L., and Mitchell-Olds, T. 2014. Natural soil microbes alter flowering phenology and the intensity of selection on flowering time in a wild Arabidopsis relative. Ecol. Lett. 17:717–726. doi:10.1111/ele.12276

96. Xu, L., Naylor, D., Dong, Z., Simmons, T., Pierroz, G., Hixson, K. K., Kim, Y. M., Zink, E. M., Engbrecht, K. M., Wang, Y., Gao, C., DeGraaf, S., Madera, M. A., Sievert, J. A., Hollingsworth, J., Birdseye, D., Scheller, H. V., Hutmacher, R., Dahlberg, J., Jansson, C., Taylor, J. W., Vogel, J. P., Lemaux, P. G., and Coleman-Derr, D. 2018. Drought delays development of the sorghum root microbiome and enriches for monoderm bacteria. Proc. Natl. Acad. Sci. USA 115:E4284–E4293. doi:10.1073/pnas.1717308115

97. Pieterse, C. M., de Jonge, R., and Berendsen, R. L. 2016. The soil-borne supremacy. Trends Plant Sci. 21:171–173. doi:10.1016/j.tplants.2016.01.018

98. Vannier, N., Agler, M., and Hacquard, S. 2019. Microbiota-mediated disease resistance in plants. PLoS Pathog. 15:e1007740. doi:10.1371/journal.ppat.1007740

99. Vorholt, J. A., Vogel, C., Carlström, C. I., and Müller, D. B. 2017. Establishing causality: opportunities of synthetic communities for plant microbiome research. Cell Host Microbe 22:142–155. doi:10.1016/j.chom.2017.07.004

100. Paredes, S. H., Gao, T., Law, T. F., Finkel, O. M., Mucyn, T., Teixeira, P. J. P. L., González-Tortuero, E., Allison, S. D., and Dangl, J. L. 2018. Design of synthetic bacterial communities for predictable plant phenotypes. PLoS Biol. 16:e2003962. doi:10.1371/journal.pbio.2003962

101. Bakker, P. A. H. M., Pieterse, C. M. J., de Jonge, R., and Berendsen, R. L. 2018. The soil-borne legacy. Cell 172:1178–1180. doi:10.1016/j.cell.2018.02.024

102. Compant, S., Samad, A., Faist, H., and Sessitsch, A. 2019. A review on the plant microbiome: ecology, functions, and emerging trends in microbial application. J. Adv. Res. 19:29–37. doi:10.1016/j.jare.2019.03.004

103. French, E., Kaplan, I., Iyer-Pascuzzi, A., Nakatsu, C. H., and Enders, L. 2021. Emerging strategies for precision microbiome management in diverse agroecosystems. Nat. Plants 7:256–267. doi:10.1038/s41477-020-00830-9

104. Reinhold-Hurek, B., Bünger, W., Burbano, C. S., Sabale, M., and Hurek, T. 2015. Roots shaping their microbiome: global hotspots for microbial activity. Annu. Rev. Phytopathol. 53:403–424. doi:10.1146/annurev-phyto-082712-102342

105. Sasse, J., Martinoia, E., and Northen, T. 2018. Feed your friends: do plant exudates shape the root microbiome? Trends Plant Sci. 23:25–41. doi:10.1016/j.tplants.2017.09.003

106. Fitzpatrick, C. R., Salas-González, I., Conway, J. M., Finkel, O. M., Gilbert, S., Russ, D., Teixeira, P. J. P. L., and Dangl, J. L. 2020. The plant microbiome: from ecology to reductionism and beyond. Annu. Rev. Microbiol. 74:81–100. doi:10.1146/annurev-micro-022620-014327

